# Imaging and quantification of prostate cancer-associated bone by polarization-sensitive optical coherence tomography

**DOI:** 10.1101/2024.02.21.581336

**Authors:** Chris Zhou, Naomi Jung, Samuel Xu, Felipe Eltit-Guersetti, Xin Lu, Qiong Wang, Doris Liang, Colm Morrissey, Eva Corey, Lawrence D. True, Rizhi Wang, Shuo Tang, Michael E. Cox

## Abstract

Prostate cancer frequently metastasizes to bone, leading to a spectrum of osteosclerotic and osteolytic lesions that cause debilitating symptoms. Accurate differentiation of bone tissue or lesion types can provide opportunity for local pathologic investigation, which is critical for understanding the bone metastasis and remains challenging. Current imaging methods lack the ability to directly differentiate tissues based on collagen organization and may induce invasive effect on bone tissues. We introduce Polarization-Sensitive Optical Coherence Tomography (PS-OCT) to investigate normal, osteosclerotic, and osteolytic bone tissues. High-resolution PS-OCT imaging reveals collagen fiber arrangement, enabling nuanced distinction of degree of collagen alignment among different bone tissue types or regions. We present a novel feature named degree of ordered organization (DOO), derived from the multiple contrasts of PS-OCT that can quantitatively evaluate bone samples from different pathologic groups, including control,osteoblastic and osteolytic tissues. The capacity of PS-OCT to differentiate trabecular/lamellar and irregular (woven bone) regions within the same specimen is tested and validated on ex-vivo samples extracted from 13 subjects. Our study is the first time that PS-OCT is applied to metastatic bone disease with the aim of enhancing the understanding of bone-related pathologies, and potentially impacting clinical practice. This work demonstrates that PS-OCT can provide useful insight into bone microstructures, and thus it has potential applications across diverse bone disorders.

## 1. Introduction

Prostate cancer (PC) is the most prevalent malignancy in men, affecting 1 in 8 men during their lifetime. Although modern therapies have dramatically increased survival, 20% of those men will advance to an incurable metastatic disease. Approximately 90% of advanced PC patients develop bone metastases (PCBM)^[REFs]^ which preferentially involve vertebrae, ribs, and epiphyses of long bones. These PCBMs give rise to debilitating symptoms, including bone pain, fractures, and spinal cord compression [3], underscoring the need for study of bone tissue samples to comprehend the intricacies of prostate cancer metastasis and develop effective treatment strategies.

PCBMs present in a broad spectrum of forms, from osteolytic to highly osteosclerotic[refs]. Although the mechanisms are still unclear, it is thought that PC cells release factors that can activate osteoblasts or osteoclasts[refs]. In healthy condition, bone resorption releases growth factors that trigger bone formation. However in PC-affected bone regions, such newly generated bone lacks regular structure which is associated with a poor mechanical performance, increasing the risk of fracture and pain. Therefore, accurate structural characterization of PCBM, and appropriate imaging of these lesions hold paramount importance in diagnosis and treatment, and in the research for better therapeutic alternatives [refs].

Conventional imaging modalities, such as X-ray [6], Computed Tomography (CT) [7] and Magnetic Resonance Imaging (MRI) [8, 9], have been widely utilized for bone tissue characterization. While micro-CT can provide excellent spatial resolution, it lacks tissue-specific differentiation. MRI, including T2-weighted imaging, excels in soft tissue contrast but may not adequately visualize the microstructures of bone [10, 11]. Quantitative back scattered electron-scanning electron microscopy (qBSE-SEM) has gained popularity for assessing bone quality and microarchitecture, offering valuable insights into trabecular and cortical bone compartments [12]. However, it relies on electron attenuation measurements and lacks penetration into deep tissue, limiting its ability in *in-vivo* measurement and clinical applications. The limited penetration also does not allow for the capture of fine (3D) details of collagen organization, which are valuable for distinguishing bone/tissue architecture and the lesion of interest. Additionally, such electron absorption counting lacks sensitivity to collagen fibers that are anisotropic, thus cannot visualize soft tissues or detect subtle changes in bone composition, particularly crucial in the context of PCBM, where soft tissue infiltration may alter bone microstructure without significant changes in bone density. Consequently, qBSE-SEM may not be sensitive enough to differentiate between subtle variations in normal bone and PCBM.

PCBM often exhibit complex interactions with the bone microenvironment [6, 13-17], leading to microstructural changes (3D spatial distribution of cells and collagen organization) that require imaging techniques with high resolution, depth penetration, and specific functionality to be effectively characterized. To address these limitations and advance bone tissue characterization, alternative imaging modalities have been explored to examine changes to mineralized tissue, including Synchrotron micro-CT [18], Raman spectroscopy [19], and X-ray diffraction [20]. However, these modalities are either highly invasive, or cannot provide depth information. Polarization-Sensitive Optical Coherence Tomography (PS-OCT), a non-invasive 3D imaging technique, with its multi-contrast sensitivity, has the potential to emerge as a powerful tool in directly differentiating between different bone tissue types [21-23]. PS-OCT leverages the polarization properties of light to unravel intricate features of bone tissue, including collagen fiber organization and lacunar morphology. This unique capability enables PS-OCT to discern nuanced differences between various bone tissue types. Moreover, PS-OCT operates at a micrometer scale resolution and millimeter scale depth penetration [23], potentially allowing for the visualization of fine 3D collagen organizations and lacunar morphology alterations associated with bone metastases.

In this study, we apply PS-OCT imaging on osteosclerotic, and osteolytic lesions generated by prostate cancer metastasis, as well as age-matched, cancer free control samples. We use qBSE-SEM to image the PCBM samples and have experts label different bone tissue types and two different bone regions (trabecular and irregular). We then develop a feature called degree of ordered organization (DOO) which combines the multiple-contrast images provided by PS-OCT. Image registration between PS-OCT and SEM is applied to locate the same region of interests (ROIs) between these two modalities. Quantification and statistical analysis are then conducted to test different bone tissue types and different bone regions. From the 13 bone samples used in this study, we found that the PS-OCT DOO image can successfully differentiate the osteoblastic tissues or the osteolytic tissues from the control group. Statistical significance is also found between the trabecular/lamellar and the irregular bone of PCBM defined on high-resolution qBSE-SEM images. By elucidating the potential clinical applications of PS-OCT in bone tissue differentiation, this work seeks to bridge the gap between cutting-edge imaging technology and clinical practice. The integration of PS-OCT into bone disorder investigation, particularly in PCBM, holds immense promise in advancing our understanding of bone-related cancer pathologies and empowering oncologists with precise and tailored therapeutic interventions.

## 2. Materials and methods

### 2.1 Sample obtention and preparation

We obtained cadaveric vertebrae samples (n=9) from 9 patients who died of metastatic PC. Patients signed written informed consent for a rapid autopsy, under the Prostate Cancer Donor Program at the University of Washington IRB#2341 [24]. We also obtained 4 vertebrae samples from 4 age-matched male control donors from the University of British Columbia Body Donation Program. These procedures, as well as the corresponding imaging process, were approved by the institutional review board at the University of Washington and the University of British Columbia (H21-02668).

Cylindrical vertebral cores were obtained using a 11 mm diameter trephine. The cylindrical specimens were transversally cut into 1/3 - 2/3 of the total length, with a water-cooled diamond saw (IsoMet 4000, Buehler, Lake Bluff, IL, USA). Out of the two bone sections (of each specimen), the smaller one was used for histology (processing methods in Section 2.2) and the larger one was prepared for SEM and PS-OCT imaging. The sections were dehydrated by using a sequence of increasing acetone concentration in ethanol (70%, 90% and 100%, repeated twice, 2 days each). The dehydrated bone sections were then infiltrated in 50%, 80% and 100% spurr embedding medium in acetone (repeated twice for each concentration, Spurr low viscosity kit, PELCO, USA) for 3 days each. After infiltration, the bone sections were embedded in spurr embedding medium. The transverse surface of the embedded bone sections was ground with a series of carbide papers and polished with 1 μm diamond suspension. The polished surface was carbon-coated and imaged in backscattered electron (BSE) mode in a scanning electron microscope (SEM, FEI Quanta 650, Oregon, USA).

### 2.2 Histological process and assessment

Bone sections obtained from the cadaveric vertebrae specimens were fixed in 10% buffered formalin for 24 hours, rinsed in phosphate-buffered saline (PBS) three times for 1 hour each, and decalcified in 10% formic acid for 5 days. The specimens were then dehydrated in an ethanol series (70%, 80%, 90%, 95%, 100%), cleared in xylene substitute, and paraffin embedded. Serial sections of 5 micrometres thickness were cut with a microtome.

The microscopy slides were stained with Masson’s trichrome, to observe the distribution of bone and other cellular components. A sample was defined as osteolytic if the bone volume was <8% with no evidence of thickened trabeculae or intertrabecular bone deposition. A sample was classified as osteosclerotic if the bone volume was >35% or if there was evidence of thickened trabeculae or bone deposited in the intertrabecular spaces [25].

### 2.3 PS-OCT system

All the samples were imaged by a custom-built Jones-matrix-based PS-OCT system (more details can be found in Ref [26]). The system is developed based on a swept-source laser (Axsun Technology Inc., MA), with the center wavelength of 1.06 μm, full width at half maximum (FWHM) bandwidth of 110 nm, and scanning rate of 100 kHz. The light from the source is split by a 90:10 single mode optical fiber coupler, where the 90% port of the coupler is connected to a passive polarization delay unit, and the 10% port to the reference arm. With a passive polarization delay unit, two orthogonal polarization states of light are separated and delayed in time. The light then passes through a 75:25 fiber coupler and the 25% port is sent to illuminate the sample. The back-scattered signal from the sample passes through the fiber coupler again and the 75% port is sent to the detection unit. Light from the sample arm and the reference arm are guided to a polarization diversity (PD) detection unit. The polarization sensitive features are measured by a Jones-matrix-based method [xx]. The average power on the sample is around 2 mW. The sensitivity of the PS-OCT system is 92 dB. The axial and lateral resolutions are measured to be ∼7.2 µm and ∼19.2 µm, respectively. Using the PS-OCT system, multiple contrasts related to the polarization of backscattered light were acquired from 3-D volumes. The horizontal scanning range can be set upon user selection between 2 mm to 12 mm. The depth detection range is usually limited by light attenuation. While the system can provide theoretical maximum depth range as ~2.8 mm (in air), the practical penetration in our practice is between 0.8 to 1.2 mm (in air). The actual penetration depth can be calculated by considering the refractive index of the bone samples. In this study, each sampling region is imaged into a 3D volumetric data of 10mm (X) by 10mm (Y) by ~1mm (Z, depth).

### 2.4 Multi-contrast imaging

By using Jones matrix detection, PS-OCT allows us to assess multiple contrasts from tissues. In this study, we utilized PS-OCT to acquire four distinct image contrasts: intensity image, phase retardation image, degree of polarization uniformity (DOPU) image, and effective signal-to-noise ratio (ESNR) image [21-23]. ESNR was also used to select the imaging depth for reliable phase retardation calculation with a threshold of 10 dB [26].

The intensity image represents the backscattered light intensity from the sample and provides a grayscale visualization of the tissue’s overall amplitude of scattering signals. It offers an anatomical overview of the bone structure, aiding in identifying different tissue regions. However, a high scattering signal may be generated from either highly organized collagen structures or highly damaged tissues, resulting in a possible ambiguity in this assessment of tissue integrity. Therefore, we do not use the intensity image for further quantification.

The phase retardation image highlights the birefringence properties of collagen fibres in bone tissue. It visualizes the phase delay of polarized light as it interacts with the tissue, offering insights into the orientation and organization of collagen fibres. When light of two orthogonal polarization states shine on aligned collagen fibres, the polarized light parallel or perpendicular to the collagen fibres generate a phase delay, which is quantified as phase retardation. When collagen fibres are randomly oriented, no phase delay is generated. Thus, strong phase retardation accumulation along the depth indicates tissue with well-aligned fibres. However, not all the well aligned fibres will exhibit strong phase retardation. For the cases when the fibres are perfectly aligned in the perpendicular orientation to the tissue surface (or parallel to the illumination beam), the phase accumulation will be low. In bone tissue, such a situation is rare and negligible.

The DOPU image quantifies the uniformity of polarization states in the tissue [27]. By applying spatial averaging on the pixel-based Stokes Vectors, DOPU can represent the randomness of spatial polarization changes. High DOPU values indicate regions with more uniform polarization states, which is related to more organized collagen micro-structures. Low DOPU values indicate regions with random polarization states (e.g. polarization scrambling), which attribute to higher level of micro-structural irregularity or higher degree of mineralization [22, 28]. Therefore, DOPU can be correlated with different bone micro-structure or mineralization changes.

The ESNR image, obtained by combining the four Jones elements, can be used to represent how subtle polarization features vary over the different polarization components [29, 30]. Basically, ESNR considers the SNR of each Jones element as well as their mutual imbalance. Strong yet randomly distributed birefringence would result in low ESNR, while moderate yet regularly organized birefringence would yield high ESNR. Therefore, ESNR can be very useful in some complex situations where bone has undergone damage and remodelling, resulting in the co-existence of both high and low birefringence.

### 2.5 Degree of ordered organization (DOO)

Several contrasts from PS-OCT can potentially be usefully in the characterization of PCBM. For example, accumulated phase retardation can indicate the organization of collagen fibres, DOPU can detect polarization scrambling caused by mineralization, and ESNR can detect polarization-sensitive changes related to fibre orientation and randomness. While each of those contrasts can indicate some aspects of PCBM, we seek to utilize all of them together to define a single parameter that provides the most comprehensive differentiation between the different bone types and bone regions. For this purpose, we have defined DOO by combining the phase retardation (φ), DOPU (Δ), and ESNR (E) as follows:

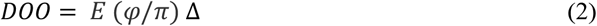

Note that the phase retardation is normalized to its maximum accumulation value, π. After combination, the DOO yields an energy-level decibel unit (dB). For each pixel, the range is set between 0 to 80 dB upon to a user selection in LabView, which is defined by examining different biology sample. In this study, the average-based en-face map results in a maximum pixel value of 58.9 dB. In this case, the maximum bound for color maping is defined as 60 dB for all the en-face DOO contrast images.

Figure 1 shows a comparison of the composite DOO contrast image with the other PS-OCT contrast images of a PCBM sample. The left column, from top to bottom, shows cross-sectional view of intensity, ESNR, phase retardation, DOPU, and DOO images, respectively. The right column shows en face view of (i) intensity, (ii) ESNR, (iii) phase retardation, (iv) DOPU, and (v) DOO images, respectively. All the en-face images in this study are obtained by using a slab of 100 pixels (~400 um) thick with average-based projection. The corresponding SEM image of the same location is shown in Fig. 1(b)-(vi) as a reference. Compared to the individual contrasts of PS-OCT, the composite DOO contrast improves the visualization of subtle features and details in the bone tissue. The arrows in Fig. 1(b)-(v) mark some example bundle-shaped features that yield high DOO values, which match very well with the highly organized bundles of collagen clusters as shown in the SEM image. Note that the high DOO regions may not necessarily be a bundle-shape. For example, there is a connective area (marked as asterisk) observed in the DOO map but not shown in the SEM image, which is obtained due to the OCT’s capability of depth imaging. This enhanced contrast of DOO facilitates the identification of microstructural alterations associated with bone metastases or other pathological conditions. Since DOO has integrated all the information from phase retardation, DOPU, and ESNR, from now on, we will mainly show the intensity and DOO images for differentiating control, osteosclerotic, and osteolytic bone tissues. All the DOO images shown in this manuscript share the same color bar, meaning that the DOO values can be compared among the DOO images from different samples. As an example, the highest DOO value comes from a control sample (Fig. 6(c) in Section 3.3).

**Figure 1.**
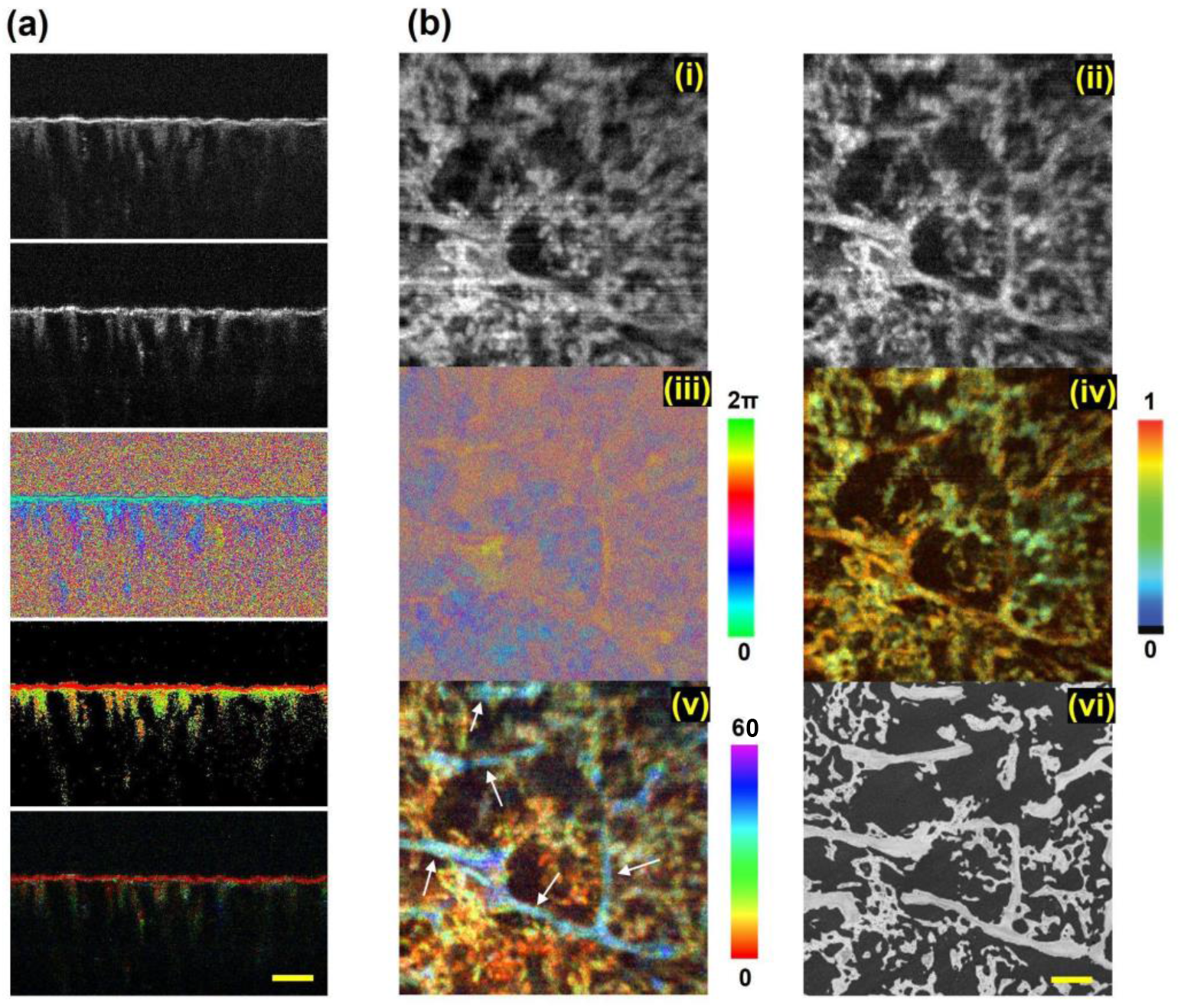
PS-OCT multiple contrast images on an example bone sample. (a) Cross-sectional images from top to bottom: intensity image, ESNR, phase retardation image, DOPU image, and DOO image. (b) ***en face*** images of the bone tissue: (i) intensity, (ii) ESNR, (iii) phase retardation, (iv) DOPU, (v) DOO, comparing the (vi) SEM image corresponding to the same location. The scale bar is 500 µm.

### 2.6 Quantification of bone regions

Figure 2 demonstrates the procedure for differencing and quantifying the two types of bone regions using qBSE-SEM and PS-OCT. Fig. 2(a) shows a high resolution qBSE-SEM image, which is used by a team of four bone experts to identify and label the trabecular and irregular bone regions, and create masks of those region using MATLAB, respectively. In principle, the qBSE images can demonstrate the organization of bone fibres in the observed 2D plane, which can to some extent represent the bone region types for certain locations. The areas with uniformly aligned/packed collagen clusters (characterized as lines and elongated bricks) are attributed as the trabecular region, while the areas with randomly distributed “woven” type collage are defined as the irregular region. Fig. 2(b) shows the mask of trabecular bone (blue) corresponding to Fig. 2(a). In Fig. 2(c), the masks generated from high resolution qBSE-SEM images are overlaid on top of the low-magnification SEM image, where the locations of the bone regions are labeled and the masks are shown as blue for trabecular and red for irregular regions. Fig. 2(d) shows the masks corresponding to Fig. 2(c). These procedures are applied to the SEM images of all samples reported in this study.

**Figure 2.**
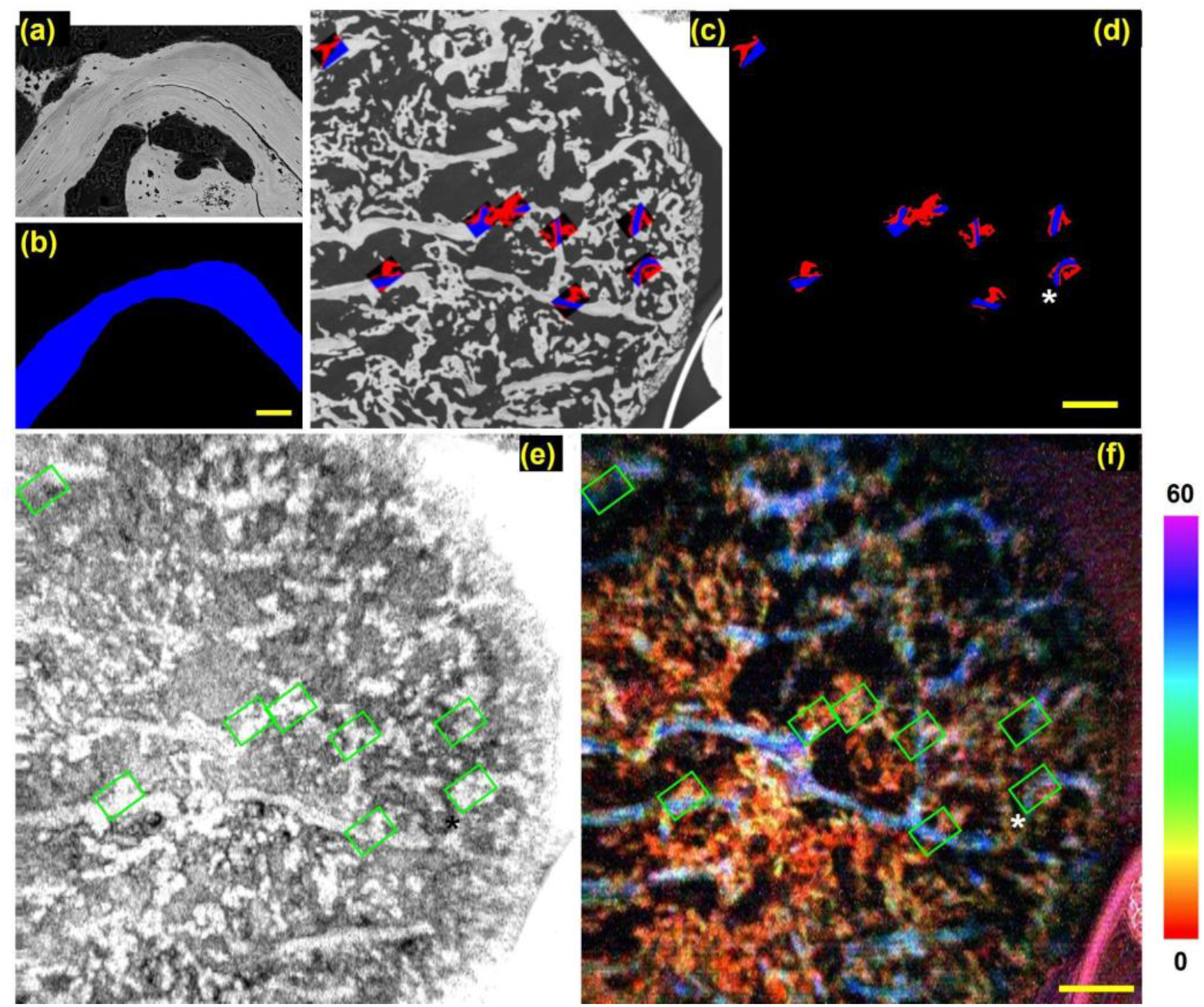
The bone sample was imaged using both qBSE-SEM and PS-OCT. The high magnification SEM image (a) was used to differentiate between the trabecular and sclerotic regions of bone. These regions were selected and converted to masks (b), and then overlaid onto the low magnification SEM image (c). By registering the low magnification SEM image with the PS-OCT image and applying the mask (d), the collagen alignment properties can be quantified for the trabecular and sclerotic bone regions using data from PS-OCT; here the locations of the quantifications are marked with green windows on the ***en face*** (e) intensity image and (f) DOO image. The blue and red region in (c) and (d) indicate the trabecular and irregular regions, respectivly. The scale bar in (a-b) is 200 μm, and in (c-f) is 1 mm.

Since the low-magnification SEM image has been adjusted and registered to the PS-OCT images, the masks generated from qBSE-SEM can now be applied onto the PS-OCT images. Figs. 2(e) and 2(f) show the ***en face*** PS-OCT intensity and DOO images, respectively, with the masks highlighted by the green boxes (the blue and red colors are not shown here in order to display the structures underneath the masks). From the PS-OCT images, the trabecular bone regions show higher scattering amplitude in the intensity image and higher degree of organization in the DOO image than irregular bone regions. Noticeably, the curvature patterns of the trabecular regions are clearly observed in the DOO images. Most of the trabecular bone regions show higher DOO values, while the irregular bone regions show lower DOO values. Therefore, the DOO image differentiates the trabecular and irregular bone regions better than the intensity image and the low-magnification SEM image. This example further highlights the utility of PS-OCT in differentiating bone regions. Afterwards, quantification methods can be applied on the PS-OCT images according to the masks.

Each PS-OCT data set can be considered as a multi-channel 3D structure. Once the 2d masks (regarded as a cluster of pixel locations) are generated, they will be attributed as segmented regions in the X-Y dimension, and then be multiplied to the 3D data set. Note that an assumption is applid here that the segemtation in X-Y plane can represent the region divisions along the sample depth. In this case, one can obtain the 3D data sets representing the trabecular region and the irregular region, respectively. A surface segmentation algorithm is applied to the data structure before segmentation to gate out all the pixel values above tissue surface. The mean and standard deviation values of each 3D region are then calculated as the quantification results. These procedures are applied to all the PS-OCT data sets (including all the possible channels) of all the samples reported in this study. For each subject, all the non-zero pixels of imaged regions are counted in the statistical analysis.**0**

## 3. Results

### 3.1 PS-OCT imaging of different bone samples

Nine PCBM samples are imaged with qBSE-SEM and PS-OCT. Figure 3 demonstrates the applicability of PS-OCT using four of these bone samples. From left to right, there are three osteoblastic samples (S1, S2, S3) and one osteolytic sample (S4). Histological images of these samples are shown in Figure 4. The type of bone sample is determined by using SEM, CT images and confirmed by histological observation of sections obtained from the same specimen (~1 mm away from SEM-PS-OCT observation plane). From the intensity ***en face*** images obtained by PS-OCT (the first row), we find that osteoblastic samples have more bone tissue compared to the osteolytic sample, which is supported by the SEM images (third row). Note that the background of sample S1 is mostly black because the overall thickness (including the embedding medium) of this sample is higher than the other samples.

**Figure 3.**
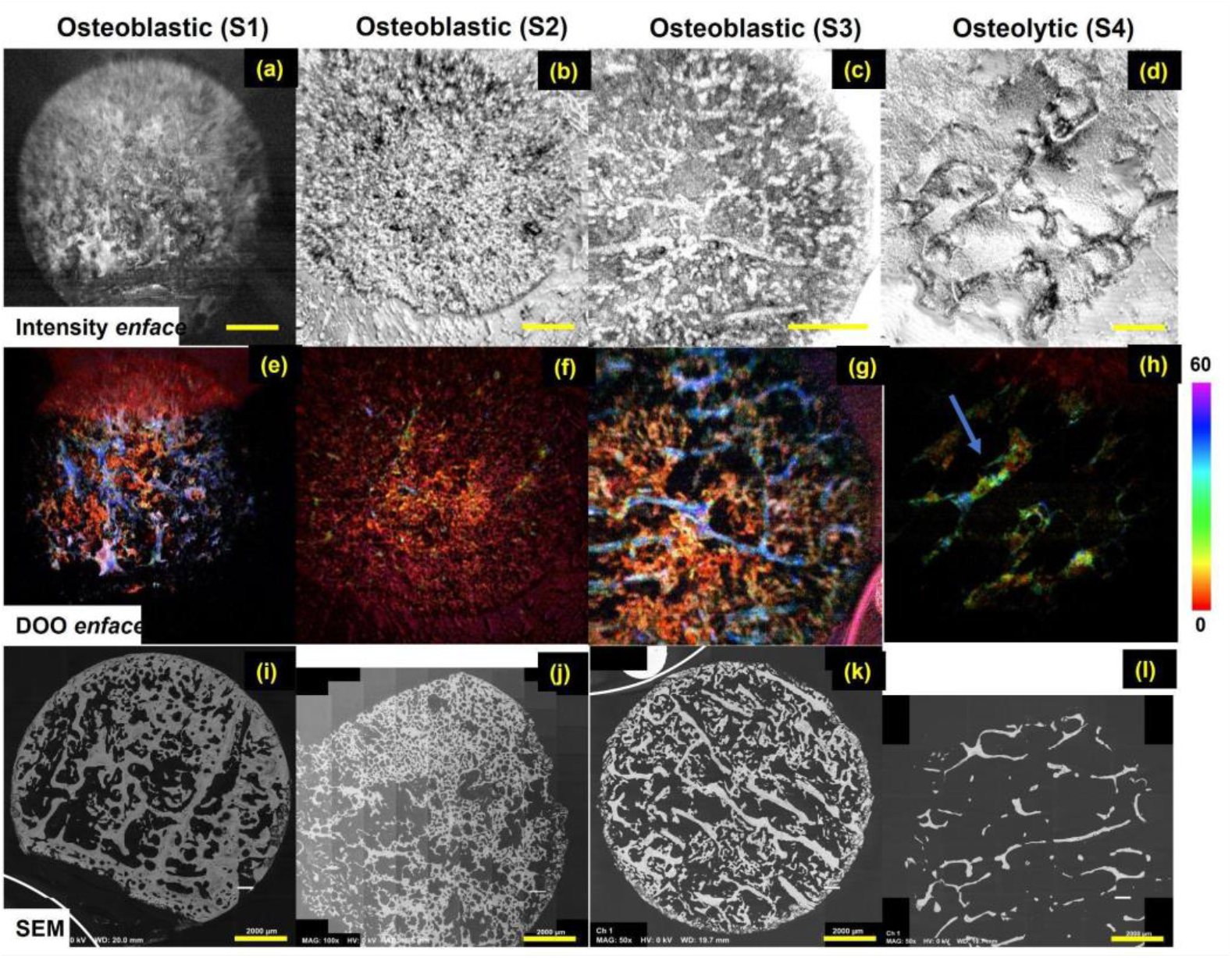
Example imaging results of three osteoblastic bone samples and one osteolytic bone sample: (a-d) PS-OCT ***en face*** intensity images, (e-h) PS-OCT ***en face*** DOO images, with corresponding (i-l) SEM images. The scale bar is 2mm, the intensity and the DOO images of each sample are co-registered and share the same scale. Possible but minor errors may exist when matching the scale of PS-OCT and the SEM images.

**Figure 4.**
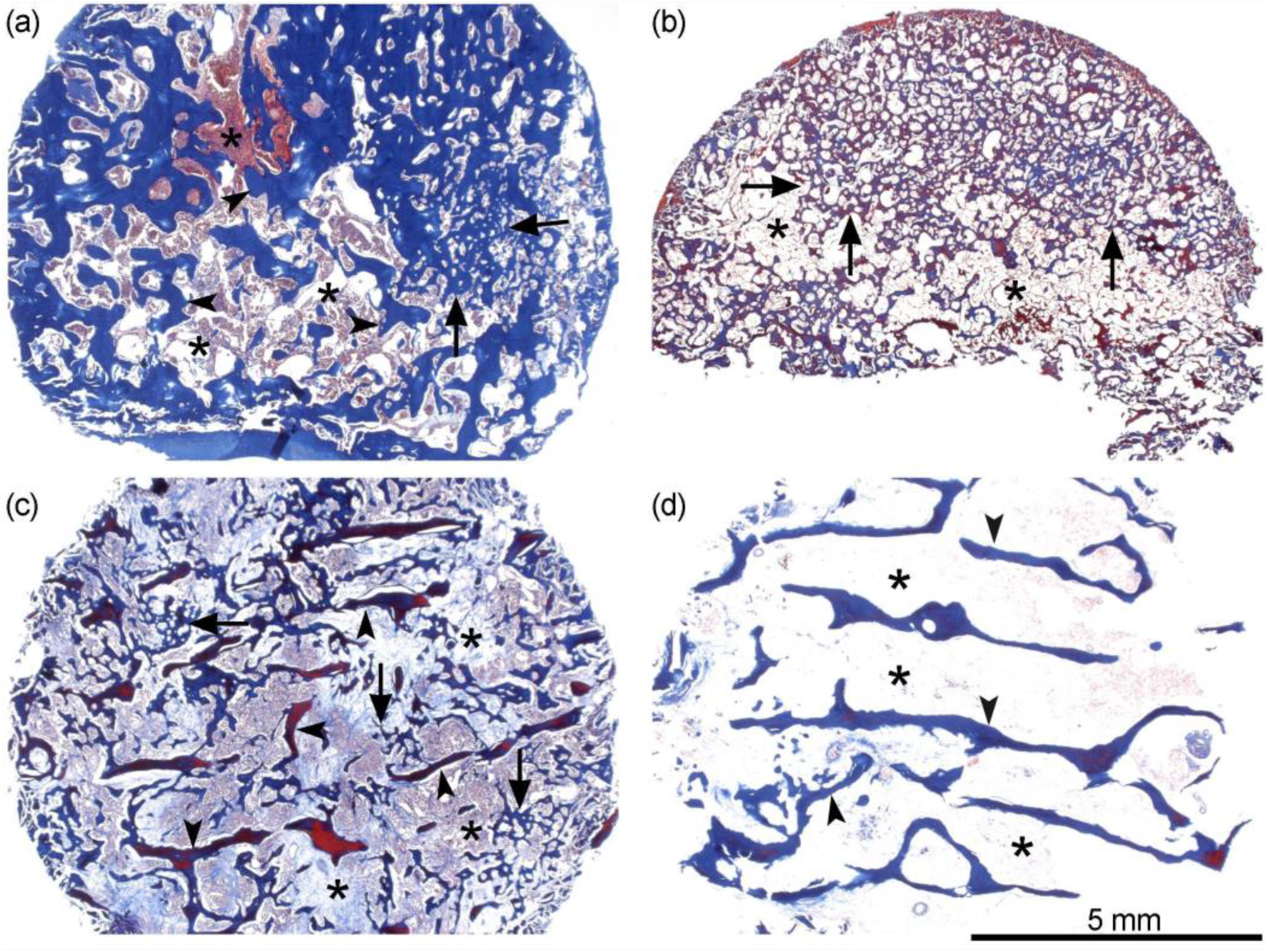
Histological images of characteristic samples analyzed with PS-OCT. (a) Sample S1, with abundant residual trabeculae (arrowheads), visible irregular prostate cancer associated bone (arrows) and reduced medullary spaces (asterisks). (b) Sample S2, with minimal if any residual trabeculae, and most of the medullary space (asterisks) occupied by irregular bone (arrows). (c) Sample S3, with evidence of trabeculae (arrowheads), and irregular bone (arrows). (d) Sample S4, osteolytic, with presence of residual trabeculae (arrowheads) this is often reduced in thickness, absence of irregular bone and enhanced medullary spaces (asterisks). Staining: Masson’s trichrome.

From the PS-OCT DOO ***en face*** images, we can observe highly organized bone features in Samples S1 and S3, but very minimally organized bone in S2. In S4, there are some residual organized bone features in the remaining bone trabeculae, however, surprisingly they are not as well aligned as the normal control samples (Fig. 6), or the organized features in the Samples S1 and S3. Such observations can be confirmed from the in-parallel SEM investigation: Sample S1 contains the most residual trabecular bone, sample S2 contains very little residual trabecular bone, and Sample S3 is the middle case (i.e. moderate amount of residual trabecular bone); Sample S4 shows some irregular bone which is not well organized. Such information is further confirmed by high resolution SEM images (e.g. Fig. 2(a)). Note that the well-organized bone in SEM images will show some textured patterns with parallel lines.Similar match up can be further confirmed by the histology images (see Figure 4).

### 3.2 Quantification and statistical results

The quantification of DOO images was then conducted on all the images captured from the three groups of samples: the normal control, osteoblastic, and osteolytic samples. Among the samples we used, the PS-OCT DOO image can be used to differentiate the osteoblastic samples (e.g. S1-S3) or the osteolytic samples (e.g. S4) from the control group. However, when accounting for all the features in all the images, the difference between the osteoblastic and osteolytic samples are statistically insignificant. This could be explained by the complexity of the bone lesions. In the osteoblastic samples, the amount of residual trabecular bone varies among individual samples. Even in the case containing a moderate amount of residual trabecular bone (e.g. S3), the features of random bone and highly organized bone coexist, which result in the broadening of the feature space. In osteolytic samples, there is a reduced amount of bone resulting from bone resorption, without any bone deposition. The DOO value of the osteolytic samples is lower than the trabecular regions in the osteoblastic samples, but higher than the irregular regions.

We then tested the difference between the trabecular/lamellar and irregular regions according to the procedures introduced in Section 2.6. For each region, we record its mean DOO value (one from lamellar and one from irregular). By combining the 72 regions over the 9 samples, we can compare the DOO value distribution and difference between the two different regions. Figure 5 shows the statistical results over the regions among the nine PCBM samples. We find that the PS-OCT DOO image is helpful to differentiate between bone regions, where we find a statistically significant difference between them (i.e. trabecular and irregular). Such a difference is validated by examining the high-resolution SEM and histology images. This result shows the possibility of using the non-invasive tool PS-OCT to identify the ROIs for specific research values/questions, without expensive (including resource, effort and time) and invasive imaging procedures (SEM, histology, etc.).

**Figure 5.**
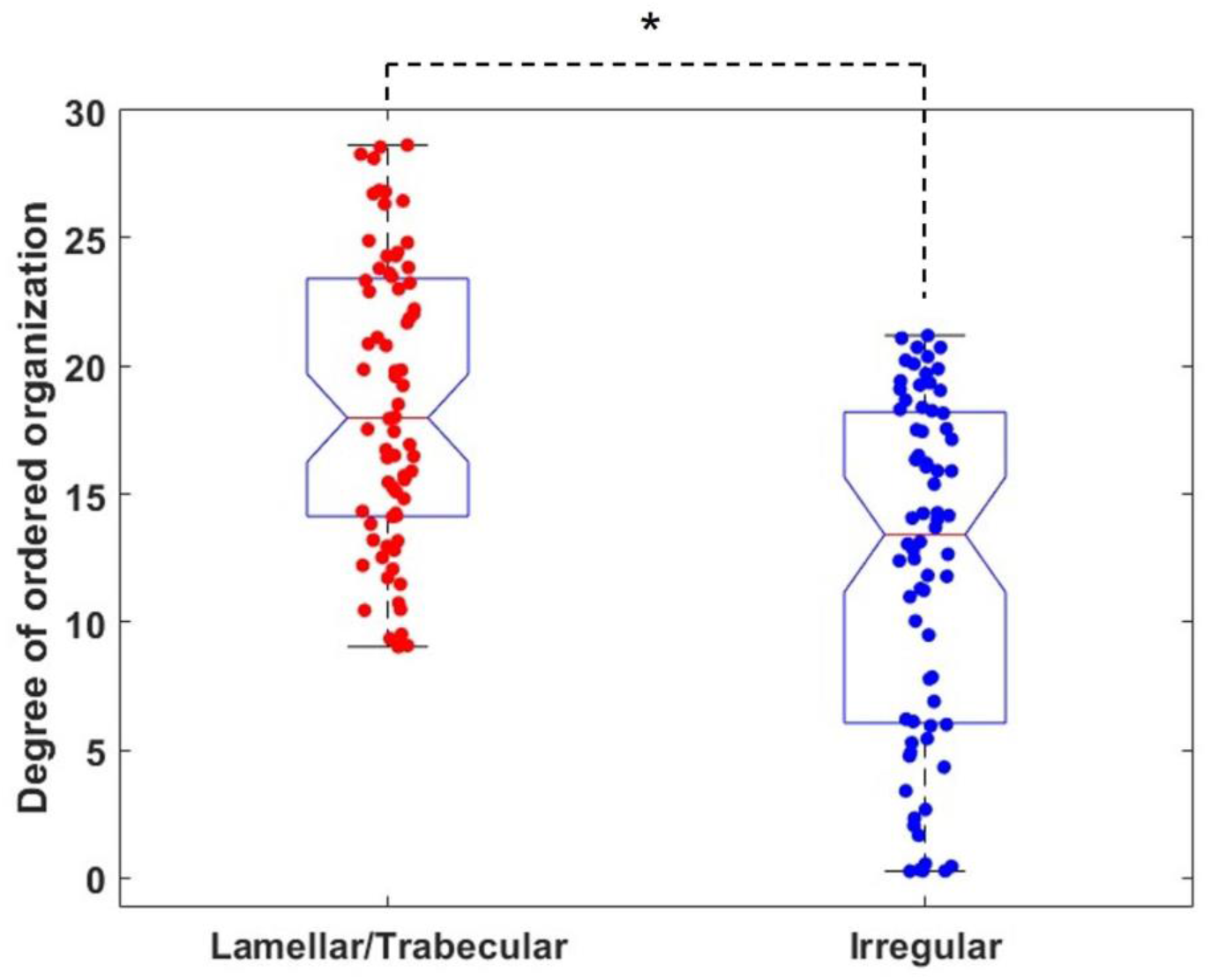
Statistical results of DOO images comparing between two bone lesion regions: lamellar and irregular. The p-value is 0.036.

### 3.3 Example data from the control group

Overall, the samples in the control group usually exhibit higher DOO compared to disease groups, which can be observed from the examples shown in Fig. 6(a-b). However, there is also sometimes a complex situation, as shown in Fig. 6(c). The example image in Fig. 6(c) actually renders very high DOO values in the middle (dark blue), surrounded by some low DOO features (red). From the images of the control samples, we confirm the diversity of the bone structures. The existence of wide distribution (both low and high DOO) and narrow distribution (the biased distribution for overwhelmingly high DOO) also shows the robustness of the quantification approach.

**Figure 6.**
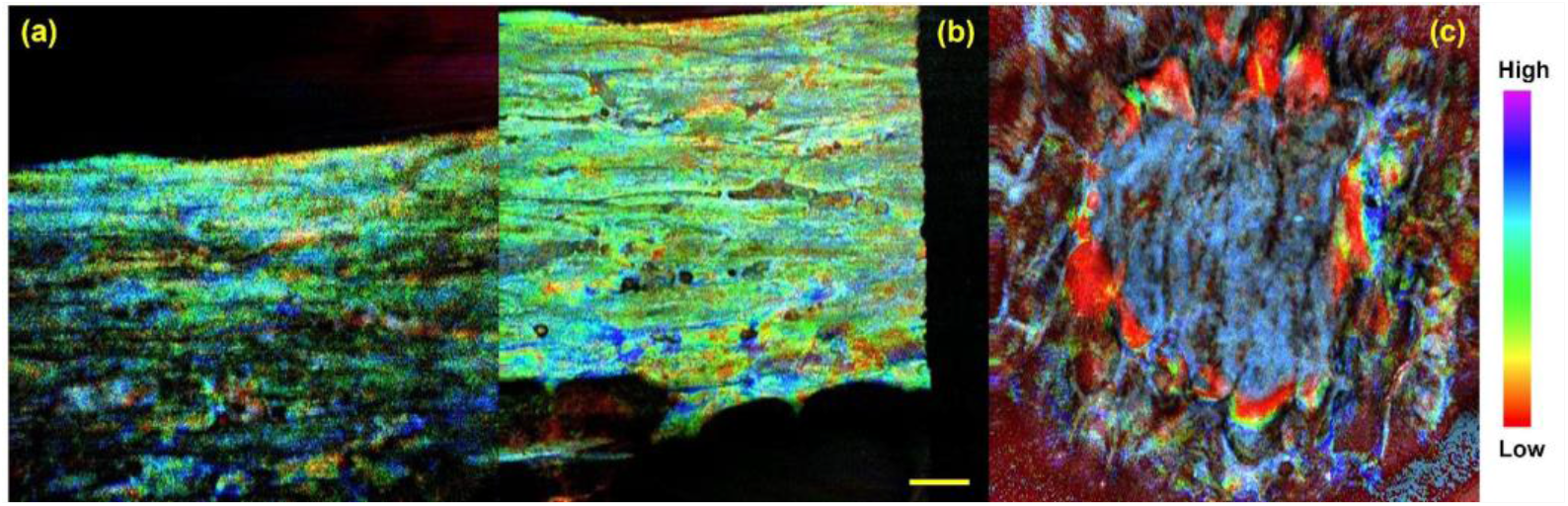
DOO images from the bone tissue samples in the control group. The scale bar is 1 mm.

**Table 1.**
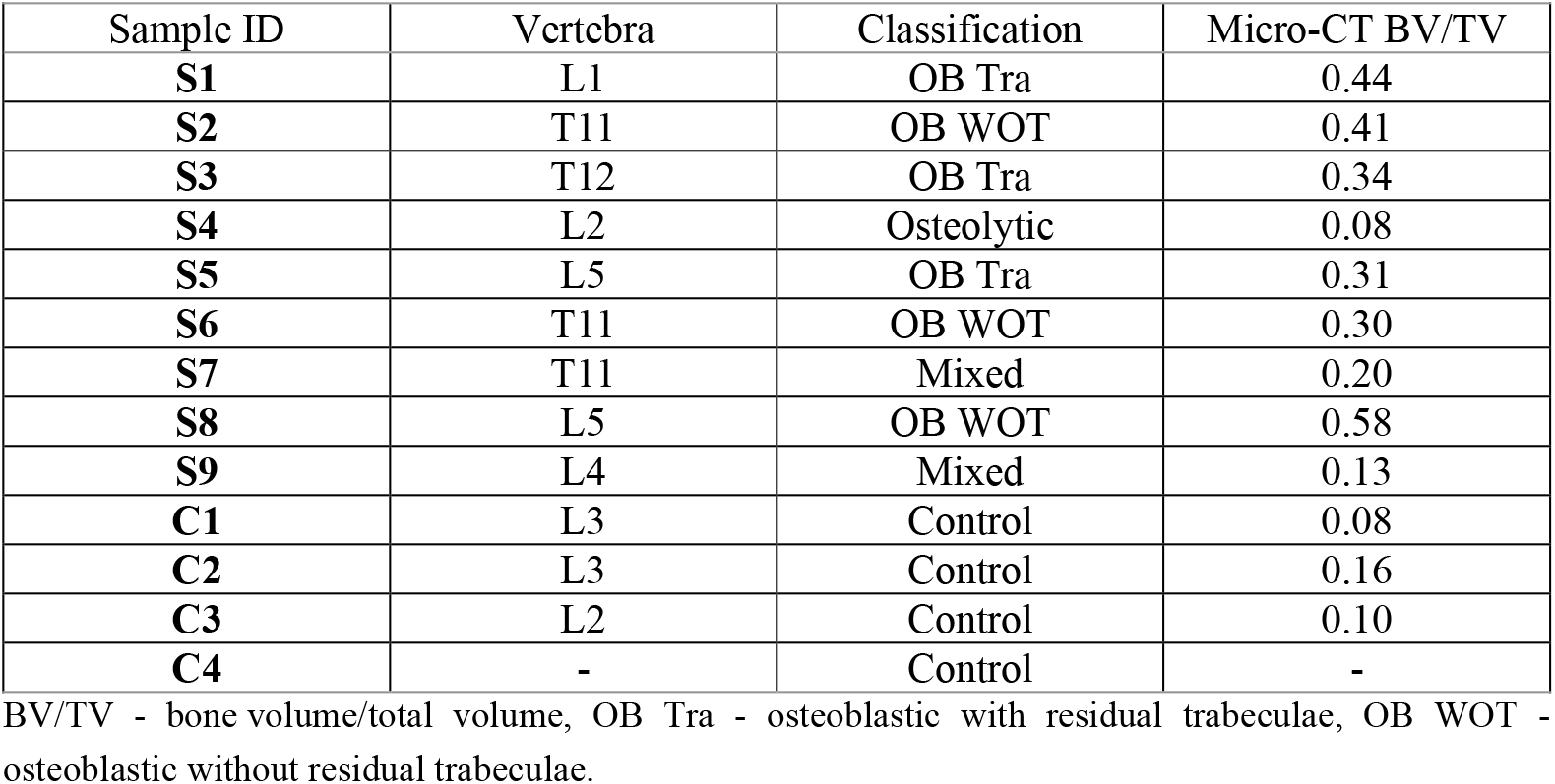
Sample data for 9 PCBM samples and 4 controls samples [24].

| Sample ID | Vertebra | Classification | Micro-CT BV/TV |
| --- | --- | --- | --- |
| **S1** | L1 | OB Tra | 0.44 |
| **S2** | T11 | OB WOT | 0.41 |
| **S3** | T12 | OB Tra | 0.34 |
| **S4** | L2 | Osteolytic | 0.08 |
| **S5** | L5 | OB Tra | 0.31 |
| **S6** | T11 | OB WOT | 0.30 |
| **S7** | T11 | Mixed | 0.20 |
| **S8** | L5 | OB WOT | 0.58 |
| **S9** | L4 | Mixed | 0.13 |
| **C1** | L3 | Control | 0.08 |
| **C2** | L3 | Control | 0.16 |
| **C3** | L2 | Control | 0.10 |
| **C4** | - | Control | - |

BV/TV - bone volume/total volume, OB Tra - osteoblastic with residual trabeculae, OB WOT - osteoblastic without residual trabeculae.

## 4. Discussions and conclusions

In this study, PS-OCT imaging is applied to different bone samples with PCBM. Exploring bone tissue characterization is pivotally significant to understanding the intricacies of PCBM and their underlying mechanisms. Using PS-OCT brought forth a remarkable revelation – an augmented DOO feature indicative of ordered collagen alignment within trabecular/lamellar bone regions. This finding may offer a glimpse into the microstructural intricacies that accompany PCBM. Importantly, DOO offers a link to the local well-aligned region as indicated in the high resolution qBSE-SEM image. Such correlative insights signify the potential of PS-OCT as a powerful tool for microstructural evaluation, which may have implications not only for PCBM but also for other bone-related disorders.

We evaluated a wide array of PCBM and age-matched controls which allowed us to decipher the intricate 3D collagen organization patterns associated with PCBM. This sheds light on the complex interplay between collagen alignment and bone tissue alterations. The insights gleaned from this approach have the potential to redefine our understanding of bone tissue dynamics in the context of PCBM. To extend this concept, we emphasize the speed of PS-OCT which can non-invasively record a 3-D volume (~250 images) within 5 seconds making it well-suited for dynamic studies.

A surprising finding in this study was the evidence of collagen disorganization in the residual trabeculae of osteolytic samples compared with residual trabeculae of osteoblastic samples and age matched controls. This differs from osteoblastic lesions in which the residual trabeculae will remain and indeed will be covered by irregular bone [Roudier 2008], and have similar organization than the control samples. These could be explained by the nature of osteolytic lesions, in which the remanent bone is in the process of being resorbed [Morrissey et al. JBMR 2013], and therefore it has altered structure observable by PS-OCT but not visible by other means. This could be a critical observation, since it suggests that the alterations in collagen structure during remodelling may start previously to the evident bone resorption process by osteoclasts, although this requires further investigation.

The integration of MATLAB-based image analysis tools, empowered by the substantial imaging depth of PS-OCT, can be used for more detailed analysis, e.g. the examination of lacunae morphology. According to the regions of high or low DOO, one can decide on more specific regions to investigate or link the collagen organization to the cellular (lacunae) quantifications. All these images can be obtained at fast speed by PS-OCT (one 3D volume within a few seconds), which will speed up the investigation and trial-and-error cycles during the research work.

There are also limitations of the current work. During the implementation of the proposed approach, we realized the hyper-sensitivity of the PS-OCT images (not only DOO) to different environmental conditions. For example, ESNR is very sensitive to the depth of observation (the intensity image is similarly affected), DOPU is very sensitive to low-scattering regions, and the phase retardation may be sensitive to residual collagen features other than from bone (such as cartilage or tendon). In processing the 2-D ***en face*** projection, surface tilting may impact the feature and cause artifacts. These detailed issues are important and need to be considered in future work.

As the PS-OCT field continues to evolve, new contrasts for this multi-contrast platform are being developed, utilizing the complexity of matrix representations of light polarization properties. For example, local birefringence can represent a similar feature as the phase retardation but offer more localized details. We have also tried to use the local birefringence in this study. The performance was similar to phase retardation when differentiating between the bone tissues (Fig. 5), where the osteoblastic and osteolytic samples are not significantly different. However, the performance (p-value) in differentiating between the two bone lesion types was actually slightly worse compared to the current case using phase retardation (Fig. 5) regardless of higher requirements for surface segmentations when using local birefringence [31, 32]. One possibility could be the actual local fibre bundles may not be as significantly different between the trabecular region and the irregular region as we expected (from this study and also the high-resolution SEM images), which can be indirectly supported from mechanical tests in our in-parallel studies. The reason that we observe such “bulk” fibre organization difference could be some group effects that require sufficient depth range to accumulate. To simplify, the “trabecular” region may not be completely trabecular but instead partially damaged and remodelled in some way that we are unable to identify from 2D imaging. To this end, optic axis (another new contrast) or more advanced depolarization metrics using Mueller matrix [33] may also be helpful to find the orientation of collagen fibers. Another reason for selecting phase retardation in this study is that we are now, at this stage, relying on a clear 2D map for understanding the bone with the support of SEM, where the depth-dependent details can be summing/averaging with detail vanishing.

Note that this study is using ex-vivo samples only. Given the nature of the complexity and difficulty of bone treatment, it is necessary to first provide some proof of concept before going to practical clinics. For in-vivo applications, we will develop a fiber-based endoscope [cite endoscope OCT] as the detection arm. Recently, Yang et al implemented endovascular OCT in imaging the spinal cord and its internal fluid environment from five animals [cite it]. From the results, the clear cerebrospinal fluid also provided an excellent medium for image acquisition, with no detectable artifact from the contents of the cerebrospinal fluid. Their work successfully proved the feasibility of spinal cord OCT imaging in living bodies, however, they did not utilize the advantages of light polarization in assessing bone features. In our future study, we will develop a similar device to conduct in-vivo vertebral imaging with PS-OCT.

In conclusion, the combination of conventional techniques (e.g. CT, SEM) with the innovative application of PS-OCT has enabled a comprehensive exploration of bone tissue alterations within PCBM. Our study demonstrates the potential of PS-OCT as an imaging tool that reveals nuanced collagen alignment variations, fostering the understanding of PCBM microstructure. This endeavour resonates not only within the domain of PCBM research but also augments our broader comprehension of bone tissue dynamics and paves the way for future advancements in bone-related pathologies.

## Acknowledgements

We thank the patients and their families, Celestia Higano, Evan Yu, Elahe Mostaghel, Heather Cheng, Pete Nelson, Bruce Montgomery, Mike Schweizer, Andrew Hsieh, Paul Lange, Jonathan Wright, Daniel Lin, Funda Vakar-Lopez, Xiaotun Zhang, Martine Roudier, Lawrence True, Robert Vessella, and the rapid autopsy teams for their contributions to the University of Washington Prostate Cancer Donor Rapid Autopsy Program. We thank the donors and families of the University of British Columbia Body Donor Program, and the contributions of Matthew Tinney, Grant Regier, Manouchehr Madani Civi, Adel Hajjay, Lexi Busse and Edwin Moore.

